# Influenza infection in ferrets with SARS-CoV-2 infection history

**DOI:** 10.1101/2022.03.22.485425

**Authors:** Caroline Vilas Boas de Melo, Florence Peters, Harry van Dijken, Stefanie Lenz, Koen van de Ven, Lisa Wijsman, Angéla Gommersbach, Tanja Schouten, Puck B. van Kasteren, van den Brand Judith, Jørgen de Jonge

**Author notes:** Address correspondence to J Jørgen de Jonge.

## Abstract

Non-pharmaceutical interventions (NPIs) to contain the SARS-CoV-2 pandemic drastically reduced human-to-human interactions, decreasing the circulation of other respiratory viruses as well. As a consequence, influenza virus circulation – normally responsible for 3-5 million hospitalizations per year globally – was significantly reduced. With downscaling the NPI countermeasures, there is a concern for increased influenza disease, particularly in individuals suffering from post-acute effects of SARS-CoV-2 infection. To investigate this possibility, we performed a sequential influenza H1N1 infection 4 weeks after an initial SARS-CoV-2 infection in the ferret model. Upon H1N1 infection, ferrets that were previously infected with SARS-CoV-2 showed an increased tendency to develop clinical symptoms compared to the control H1N1 infected animals. Histopathological analysis indicated only a slight increase for type II pneumocyte hyperplasia and bronchitis. The effects of the sequential infection thus appeared minor. However, ferrets were infected with B.1.351-SARS-CoV-2, the beta variant of concern, which replicated poorly in our model. The histopathology of the respiratory organs was mostly resolved 4 weeks after SARS-CoV-2 infection, with only reminiscent histopathological features in the upper respiratory tract. Nevertheless, SARS-CoV-2 specific cellular and humoral responses were observed, confirming an established infection. Thus, there may likely be a SARS-CoV-2 variant-dependent effect on the severity of disease upon a sequential influenza infection as we observed mild effects upon a mild infection. It, however, remains to be determined what the impact is of more virulent SARS-CoV-2 variants.

**Importance:** During the COVID-19 pandemic, the use of face masks, social distancing and isolation were not only effective in decreasing the circulation of SARS-CoV-2, but also in reducing other respiratory viruses such as influenza. With less restrictions, influenza is slowly returning. In the meantime, people still suffering from long-COVID, could be more vulnerable to an influenza virus infection and develop more severe influenza disease. This study provides directions to the effect of a previous SARS-CoV-2 exposure on influenza disease severity in the ferret model. This model is highly valuable to test sequential infections under controlled settings for translation to humans. We could not induce clear long-term COVID-19 effects as SARS-CoV-2 infection in ferrets was mild. However, we still observed a slight increase in influenza disease severity compared to ferrets that had not encountered SARS-CoV-2 before. It may therefore be advisable to include long-COVID patients as a risk group for influenza vaccination.

## Introduction

Two years after the emergence of SARS-CoV-2 in late 2019, the causative agent of COVID- 19 is still an important global health and economic problem. In order to mitigate virus transmission, a set of non-pharmaceutical interventions (NPIs), such as the use of face masks, physical distancing and travel restrictions were adopted worldwide, leading to a considerable decrease in social interactions [1]. The unprecedented reduced human interaction was not only effective against SARS-CoV-2 transmission, but it also resulted in an almost complete absence of circulation of several other respiratory viruses [2–4]. For influenza virus, for example, the recorded peak of over 40.000 weekly cases in the global 2019-2020 influenza season, dramatically decreased in incidence from March 2020 to essentially zero in the following months, leading to the absence of the global 2020-2021 influenza season [5]. With the release of NPIs and subsequent renewed circulation of respiratory viruses, there is a concern that influenza infection might lead to more severe disease in individuals previously exposed to SARS-CoV-2. There are at least two reasons for this: (1) the interrupted seasonal exposure has resulted in reduced immunity in the population, which is therefore more susceptible [6]; and (2), residual effects of COVID-19 can have a negative clinical impact on a sequential respiratory viral infection, such as influenza.

Fatigue, dyspnea, and chest pain are commonly reported after cleared SARS-CoV-2 infection [7]. These long-term effects of COVID-19, so-called post-acute COVID-19 or long-COVID syndrome are possible clinical repercussions of pulmonary sequelae of a SARS-CoV-2 infection [8, 9]. The only evidence for increased disease in a situation where SARS-CoV-2 and influenza co-circulated goes back to the beginning of the SARS-CoV-2 pandemic, when influenza virus was still circulating. It was then observed that a higher risk for severe outcome was associated with influenza co-infection in lethal cases of COVID-19 [10–13]. In the current stage, however, it is still unclear whether a recent SARS-CoV-2 infection history could negatively impact a sequential influenza infection.

Animal models are an ideal tool to investigate combinations of infections under controlled conditions. Multiple animal models of COVID-19 are currently available [14], but not all of them are equally suitable for modelling influenza disease. So far, co-infection studies with SARS-CoV-2 and influenza A virus (IAV) in hamster and K18-ACE2 mouse models indicate more severe pneumonia [15–17] and confirm the initial findings in humans [18]. Ferrets co-infected with SARS-CoV-2 and IAV developed increased weight loss and enhanced inflammation in the nasal cavity and lungs [19]. In transgenic hACE2 mice, more severe lung damage upon a sequential infection with H1N1 with a brief time interval (7 dpi) and during the convalescent SARS-CoV-2 infection phase (14 dpi) [20] was observed. Little is known, however, about the effects of an influenza infection following a resolved SARS-CoV-2 infection or during post-acute COVID-19, which is the more likely scenario.

Ferrets are the best small animal model for influenza infection and disease and they are also susceptible for SARS-CoV-2 infection, albeit reproducing only mild or non-clinical COVID- 19 [14, 21, 22]. Nevertheless, follicular hyperplasia in the higher airways of the lung and inflammatory infiltration in the nasal cavity has been observed 21 days after experimental infection with SARS-CoV-2, when the viral infection was long cleared [23]. These observations could reflect, in part, the long-term disease (post-acute COVID-19) characterized by prolonged respiratory complaints and fatigue. We experimentally infected ferrets with SARS-CoV-2 beta variant (B.1.351), a variant of concern (VOC) at the time these experiments were performed. The ferrets were followed up to 4 weeks post infection, with the goal to resemble post-acute COVID-19 and were then infected with an H1N1 influenza virus. Our aim was to investigate the impact of the previous exposure to SARS-CoV-2 on the severity of disease and pathology of respiratory organs upon a sequential influenza (H1N1) infection, and to provide evidence to support influenza vaccination as a strategy for mitigating the said severe effects.

## Methods

### Ethics statement

All animal procedures were conducted in accordance to the European regulation for animal experimentation. The study proposal was evaluated and approved by the Animal Welfare Body of Poonawalla Science Park – Animal Research Center (Bilthoven, The Netherlands) under permit number AVD32600 2018 4765 of the Dutch Central Committee for Animal experiments.

### Ferrets

Twenty outbred male ferrets (*Mustela putorius furo*), aged 8 months, were obtained from the colony of Euroferret (Denmark). The ferrets were investigated for previous Aleutian disease, canine herpes virus and ferret coronaviruses infections by measuring antibody titers in serum (supplementary table 1). All ferrets tested negative for antibody responses measured against SARS-CoV-2 spike protein (S), its receptor binding domain (RBD), and the nuclear protein of influenza virus by ELISA. The ferrets were semi-randomly allocated in three matching groups by weight, with an average of 1.3 kg ± 0.1 kg, housed in closed cages and acclimated for 24 days in the designated experimental groups.

### Cell culture and virus isolates

SARS-CoV-2 beta variant (B.1.351) was isolated from a COVID-19 patient (hCoV- 19/Netherlands/NH-RIVM-20159/2021, nextclade 20H/501Y.V2). Influenza A virus (H1N1) A/Michigan/45/2015 was obtained from National Institute for Biological Standards and Control (NIBSC, London, England - code: 16/354). SARS-CoV-2 alpha variant (B.1.1.7- 484K) was isolated from a COVID-19 patient (hCoV-19/Netherlands/UT-RIVM-12844/2021). SARS-CoV-2 virus culture was performed in VERO E6 cells (Vero C1008, ATCC CRL-1586), first grown in Dulbecco’s Modified Eagle Medium (DMEM, Gibco) supplemented with 10% of fetal bovine serum (FBS) + 1x penicillin, streptomycin and glutamine (PSG) at 37°C and 5% CO_2_. At a confluency of 90-95%, VERO E6 cells were washed twice with PBS and a virus suspension of SARS-CoV-2 in infection medium (DMEM + 1x PSG) was added to the cells and incubated at 37°C and 5% CO_2_ for 2-3 days. When a cytopathological effect (CPE) of 90% was observed, the culture supernatant was collected and spun down for 10 min at 1000xg a room temperature (RT) to harvest the virus suspension. Wild-type mumps virus (MuVi/Utrecht.NLD/40.10; genotype G) was cultured on VERO cells in infection medium containing DMEM + 1x PSG + 2% FBS and the virus was harvested when a CPE of >90% was observed. MDCK cells were grown in Minimum Essential Medium (MEM, Gibco) + 10% FBS + 1x PSG at 37°C and 5% CO_2_ until a confluence of ≥90%. MDCK cells were washed twice with PBS, and H1N1 virus suspension was added to the cells in MEM + 1x PSG + 2 µg/ml L- 1-tosylamido-2-phenylethyl chloromethyl ketone (TPCK) treated trypsin. The virus culture was incubated at 37°C and 5% CO_2_ for 2-3 days, and the virus suspension was harvested when a CPE of 90% was observed. Aliquots of each of the virus cultures were snap frozen and stored at -80°C, and viral titers were determined by TCID_50_ assay on VERO E6 cells for SARS-CoV- 2 or MDCK cells for H1N1. The SARS-CoV-2 beta variant virus stock was sequenced and did not contain mutations compared to the reference isolate. The SARS-CoV-2 alpha variant 484K acquired cell culture-induced mutations in amino acids within the Open Reading Frame (ORF) regions: two deletions in ORF1a:V84- and ORF1a:M85- and one substitution in ORF1a:L3829F. Due to the mutations, the alpha variant was used only for ELISpot stimulations where it is not expected to interfere with the experiments.

### Animal handling and sample collection

Ferrets were anesthetized with ketamine (intramuscular, 5mg/kg) for weight measurements and swab collection. For drawing blood, infection and surgical handling, medetomidine (0.1mg/kg) was added in combination with ketamine and antagonized with antisedan (0.25mg/kg). Pre-analgetic was administered subcutaneously (Carporal, 0.2mL) prior to implantation of transponder for body temperature measurements (Star Oddi, Iceland) in the peritoneal cavity of the ferrets 20 days before start of the experiment. Nose and throat swabs were collected in tubes containing viral transport medium (15% sucrose, 2.5 μg/ml Amphotericin B [Merck], 100 U/mL penicillin, 100 μg/mL streptomycin and 250 μg/mL gentamicin [Sigma]), vortexed and stored at -80°C for later analysis. During the experiment, blood was collected from the cranial vena cava in vacutainer tubes (BD) coated with heparin for PBMC isolation, or in vacutainer tubes (BD) containing polymer to obtain serum. Euthanasia was performed by exsanguination via heart puncture under anesthesia with ketamine and medetomidine. At necropsy, nasal turbinates, trachea and lungs were collected for pathological and virological analysis. The lower portion of the trachea, right nasal turbinates and three representative lung sections were collected from the right cranial, middle and caudal lobe in Lysing Matrix A tubes (MP Biomedicals, Germany) for virology assays. The trachea and lung samples for virology were stored at -80°C until processed for tissue dissociation in infection medium (DMEM + 2% FBS + 1x PSG). Animals and samples collected up to 9 days after SARS-CoV-2 infection were handled under BSL-3 conditions. After testing negative for infectious SARS-CoV-2 virus, the samples were handled under BSL-2 conditions. All dissections were performed under BSL-3 conditions. H1N1-infected samples were handled under BSL-2 conditions.

### Study design and infections

Twelve ferrets were infected intranasally (i.n.) with 10_7_ TCID_50_ of SARS-CoV-2 beta VOC diluted in phosphate buffered saline (PBS) on day 0. An inoculum volume of 1 mL was used, with the aim to reach the lower regions of the respiratory tract. Six out of twelve ferrets were euthanized at 28 days post infection (d.p.i.) to describe the post-acute phase of SARS-CoV-2 beta VOC infection in ferrets. The remaining six SARS-CoV-2 infected ferrets were infected i.n. with 10_6_ TCID_50_ of H1N1 diluted in PBS. A control group of five ferrets infected only with H1N1 was included. For capacity reasons, H1N1 infections were performed on two separate days: on 29 d.p.i. (experiment A) and on 30 d.p.i. (experiment B) and followed up to 5 days post H1N1 infection. The results of experiments A and B are merged. The mock-infected group of ferrets (n=3) received 1 mL of PBS i.n. only, on the days of each different infection (SARS-CoV-2 on day 0, H1N1 on days 29/30) and were followed up to the end of the experiment (35 d.p.i.). Nose and throat swabs for virology assays were collected on days 0, 3, 5 and 9 for SARS-CoV-2 infection and on days 29 to 35 for H1N1 infection (0, 2, 4 and 5 days after H1N1 infection). Blood was obtained from the cranial vena cava on days 0, 14, 21 and 29 during the study, and by heart puncture on the days of euthanasia (28 and 35 d.p.i.) (figure 1a).

**Figure 1.**
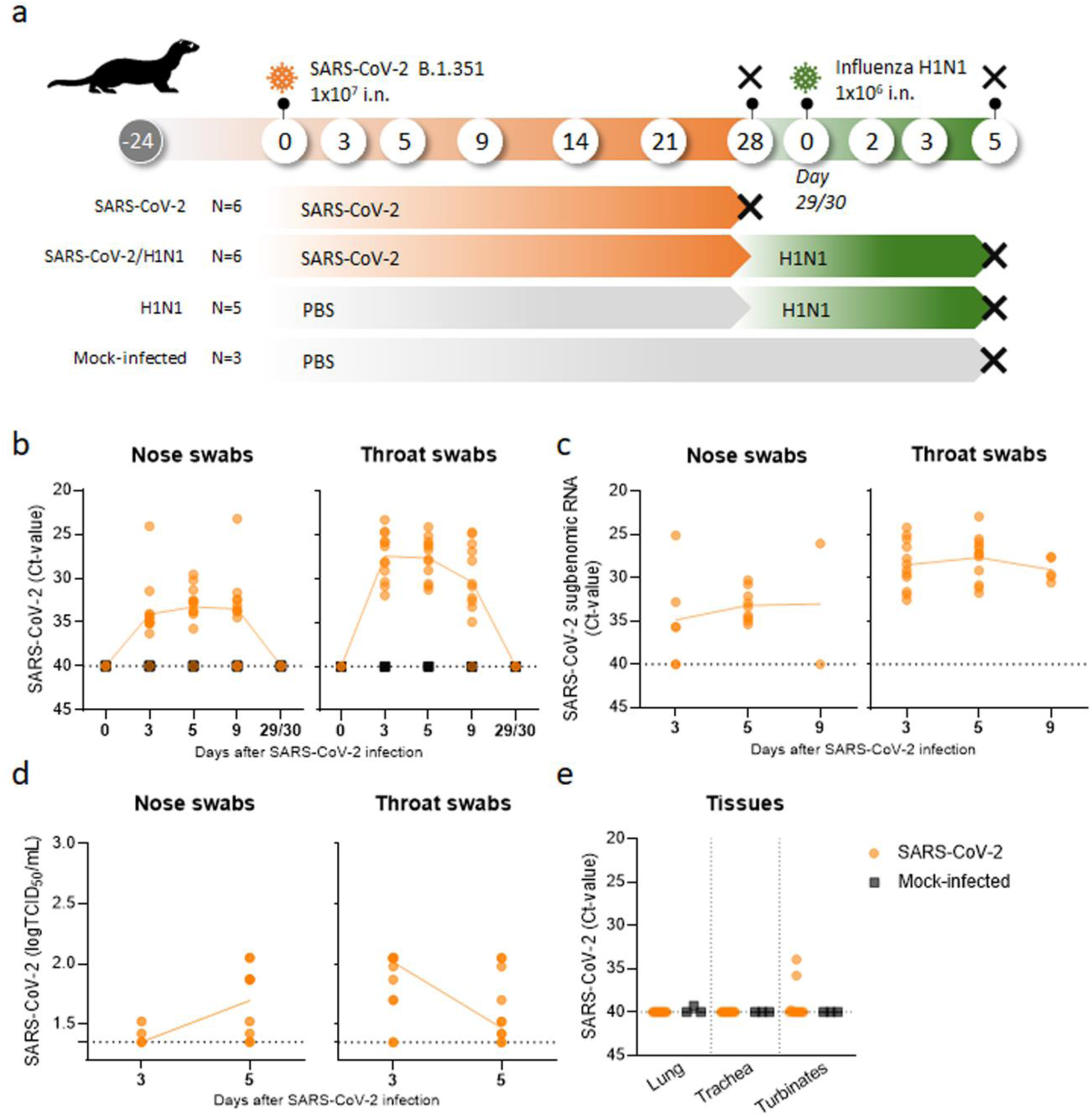
Study design and SARS-CoV-2 (B.1.351, beta VOC) viral load. **(a)** Male ferrets were infected intranasally (i.n) with SARS-CoV-2 (B.1.351, beta VOC), H1N1, or with SARS-CoV-2 and H1N1 in sequence. Control groups were injected i.n. with phosphate buffered saline (PBS, mock-infected). Two groups of six ferrets each were infected with beta VOC on Day 0. One of these groups was followed up to 28 days post-infection (d.p.i.) and then euthanized (black cross). The other group was sequentially infected with H1N1 on days 29/30. One group of five ferrets was single-infected with H1N1 at the same timepoint. H1N1- and mock-infected ferrets were followed up to five days after H1N1 infection and then euthanized. **(b-e)** Viral load and virus replication of all beta VOC-infected ferrets (n=12, orange dots) and mock-infected ferrets (n=3, black squares). **(b)** Beta VOC viral RNA detected 0, 3, 5, 9 and 29 d.p.i. Only the remaining six beta VOC-infected ferrets were tested to ensure the absence of the virus at the moment of receiving the H1N1 infection (day 29/30). Beta VOC subgenomic RNA detected by RT-PCR **(c)** and viral replication detected by TCID_50_ assay **(d)** in nose and throat swabs 3 and 5 d.p.i. **(e)** Beta VOC viral RNA in lung, trachea and nasal turbinates 28 d.p.i. by RT-PCR. Horizontal dotted lines depict the limit of detection by RT-qPCR, set at a Ct-value of 40 **(b, c and e)** and 1.3 log TCID_50_/mL for TCID_50_ assay. Connecting lines depicts mean and dots represent individual observations.

### Clinical evaluation

Clinical examination was performed daily, for nine days post SARS-CoV-2 infection and for five days post H1N1 infection. A score for clinical signs was employed relative to other ferret groups. Clinical signs comprise decreased behavioral activity, increased severity of breathing, nasal discharge and sneezing. The clinical endpoints were based on activity (0 = active; 1 = active when stimulated; 2 = inactive and 3 = lethargic) and breathing (0 = normal; 1 = fast breathing; 2 = heavy/stomach breathing); ferrets were euthanized if lethargic or presenting a combined score of 4 for activity and breathing [23]. Body temperature was measured by the implanted temperature transponder every 30 minutes from five days prior to each infection. Fever was calculated as temperature deviations from the baseline temperature, measured during the five days prior to infection. Variations in body weight were determined relative to the baseline body weight on day 0 of each infection.

### Viral load by RT-qPCR

Swabs and tissues samples for RT-qPCR were dissociated in lysis buffer containing equine arteritis virus (EAV) as an internal RT-qPCR control. Extraction of total nucleic acid was performed using the MagNA Pure 96 DNA and Viral NA Small Volume Kit in the MagNA Pure 96 system (Roche, Penzberg, Germany), and eluted in a volume of 50μl Roche Tris-HCl elution buffer. A 20μl Real-time Reverse-Transcription PCR (RT-qPCR) reaction contained 5μl of sample nucleic acid, 7μl of 4x Taqman Fast Virus Master Mix (Thermo Fisher), 5μl of DNAse/RNAse free water and 3μl of primers and probe mix were used. SARS-CoV-2 was detected using the E-gene forward (ACAGGTACGTTAATAGTTAATAGCGT) and reverse (ATATTGCAGCAGTACGCACACA) primers and probe (ACACTAGCCATCCTTACTGCGCTTCG). EAV was detected using the E-gene forward (CTGTCGCTTGTGCTCAATTTAC) and reverse (AGCGTCCGAAGCATCTC) primers and probe (TGCAGCTTATGTTCCTTGCACTGTGTTC). For H1N1 the primers and probe are designed for the H1N1pmd09 and was detected using the E-gene forward (TGGACTTACAATGCCGAACT) and reverse (CAGCGGTTTCCAATTTCCTT) and probe (GGACTATCACGATTCAAATGTGAAGAACT). RT-qPCR was performed for 10 minutes at 95°C, followed by 40 cycles of 95°C for 30 seconds and primer-specific annealing temperature of 60°C for 45 seconds, as previously described [23–25]. This was followed by 95°C for 15 seconds, 53°C for 60 seconds and 95°C for 15 seconds. Subgenomic mRNA of SARS-CoV-2 was detected by in-house subgenomic mRNA E-gene assay, using the Egene reverse primer and probe and the forward primer as previously described [26]. All experiments were performed on a Light Cycler 480 I (LC480 I, Roche). Cycle threshold (Ct) values were recorded for genomic and subgenomic RNA, to determine viral burden and infectious virus, respectively.

### TCID_50_ determination

SARS-CoV-2 virus stock, swab material and tissue samples were 1:10 serial diluted in DMEM medium containing 2% FBS and 1x PSG and added on VERO E6 cells in 96-well plates. H1N1 virus stock, swabs material and tissue samples were 1:10 serial diluted MEM medium + 1x PSG + TPCK treated trypsin 2µg/ml and added on MDCK cells in 96-well plates. Virus stocks were titrated in octuplicate, while swab material and tissue samples were titrated in sextuplicate. The plates were incubated at 37°C and CPE was scored after 5 days (H1N1) or 6 days (SARS-CoV-2). TCID_50_ values were calculated using the Reed & Muench method.

### Histopathology

Tissues harvested for histological examination (trachea, bronchus and left lung) were fixed in 10% neutral-buffered formalin, embedded in paraffin, sectioned at 4 µm and stained with hematoxylin and eosin (HE) for examination by light microscopy. Histopathology was scored blindly by a veterinary pathologist. Semiquantitative assessment of influenza virus-associated inflammation in the lung was performed on a longitudinal- and cross-section of the cranial and caudal lobe as reported earlier [27] with few modifications. Briefly, for the extent of alveolitis and alveolar damage we used: 0, 1 (1-25%), 2 (25-50%) or 3 (>50%). For the severity of alveolitis, bronchiolitis, bronchitis, and tracheitis we scored: 0, no inflammatory cells; 1, few inflammatory cells; 2, moderate numbers of inflammatory cells; 3, many inflammatory cells. For the presence of alveolar oedema and type II pneumocyte hyperplasia we scored: 0, 1 (1- 25%), 2 (25-50%) or 3 (>50%). The presence of alveolar hemorrhage we scored: 0, no; 1, yes. For the extent of peribronchial/perivascular oedema we scored: 0, no, 1, yes. Finally, for the extent of peribronchial, peribronchiolar, and perivascular infiltrates we scored: 0, none; 1, one to two cells thick; 2, three to ten cells thick; 3, more than ten cells thick. The average scores for size and severity of inflammation of the different slides provided the total score per animal.

### Antibody responses

Antibody responses were measured against recombinant beta-SARS-CoV-2 B.1.351 spike protein (S) and its receptor binding domain (RBD) (10777-CV and 10735-CV, R&D Systems) by ELISA as previously described [23]. Briefly, ferret serum was pre-diluted 1:50 in PBS + 0,1% Tween 80 solution (dilution buffer) and ELISA assay was performed in 2-fold serial dilutions to determine the optimal fitted concentration curve. Next, the plates were incubated with HRP-321 conjugated goat anti-ferret IgG (ab112770, Abcam), diluted 1:5000 in dilution buffer + 0.5% powdered milk. Color development was obtained by incubation with KPL Sure Blue TMB Microwell Peroxidase Substrate 1-Component (95059-286, VWR), and the reaction was stopped using 2M H_2_SO_4_ (80012010.2500, Boom). Antibody concentrations were measured by optical density (O.D.) at 450 nm absorbance using an EL808 absorbance reader (Bio-Tek Instruments) and represented at a 1:100 or 1:200 sera dilution for Spike protein and RBD, respectively, over the different study timepoints.

### PBMC isolation

Peripheral blood mononuclear cells (PBMC) were isolated from 1:1 PBS-diluted blood by density gradient centrifugation (1:1 LymphoPrep 1114547 and Lympholyte-M CL5035, Sanbio). The cells were isolated from the interphase after centrifugation at 800xg. After two washing steps in RPMI 1640 + 1% FBS, the final cell pellet was resuspended in stimulation medium (RPMI1640 + 10% FBS + 1x PSG) at two concentrations: 2,5 × 10_5_ cells/mL for ELISpot assays and 3 × 10_6_/mL for flow cytometry assay.

### ELISpot

Pools consisting of 15-mer overlapping (11 amino acids) peptides of entire SARS-CoV-2 proteins from the original SARS-CoV-2 strain (1 µg/ml, PepMix, JPT), were used for PBMC stimulation. For the spike protein, the peptides are divided over two vials with complementary sequences: 1-158 and 159-315. PBMCs were also stimulated with live alpha-SARS-CoV-2 (B.1.1.7) or beta-SARS-CoV-2 (B.1.351) at 2.5 × 10_4_ TCID_50_ (MOI 0.1). Mumps virus (2.5 × 10_5_ TCID_50_, MOI 1) and HIV peptide pool (1µg/ml, PepMix, JPT) were included as negative controls for live-virus and peptide pool stimulations, respectively. Staphylococcal Enterotoxin B (SEB) was used as superantigen for lymphocyte activation and served as a positive control for T cell stimulation. Medium-stimulated condition was included to correct for background activation. The cells were incubated for 20 hours at 37°C in 5% CO_2_ incubator in pre-coated Ferret IFNγ-ELISpot plates (3112-4APW-2, Mabtech). The assay was performed following the manufacturers protocol, with one modification in the incubation time with the primary antibody to overnight at 4°C. The plates were air-dried for at least 2 days and further heat-treated at 65°C for 3 hours to ensure inactivity of SARS-CoV-2 prior to scan and analysis using ImmunoSpot S6 CORE (CTL, Cleveland, OH). Background was corrected by subtraction of medium-stimulated spot counts from every other stimulation.

### IFNγ flow cytometry

PBMCs at a concentration of 3 × 10_6_ cells/mL were stimulated with indicated live virus for 20 hours and peptide pools for 6 hours at 37°C in 5% CO_2_ incubator. In the last 5 hours, Brefeldin A solution (Biolegend) was added to the stimulated PBMCs to prevent secretion of intracellular cytokines. The stimulated cells were spun down at 500xg/3min and resuspended in 150µl of washing buffer containing PBS + 0.5% bovine serum albumin + 2 mM EDTA. The cell suspension was transferred into a 96-well plate for flow cytometry. Extracellular staining was performed using APC-conjugated anti-CD4 (60003-MM02-A, clone 02, Sino Biological), eFluor450-conjugated anti-CD8a (17008642, clone OKT8, eBioscience) and FVS780- conjugated cell viability staining for 30 minutes/4°C. The cells were washed twice and intracellular staining was performed using the FoxP3 staining kit (eBioscience). Fixation of the cells occurred with incubation of Fixation/Permeabilization buffer for 20 min/4°C, followed by two washing steps with 1x permbuffer, prior to incubation for 30 min/RT with the antibodies: FITC-conjugated anti-CD3e (MCA1477A647, clone CD3-12, BioRad) and PE-conjugated anti-IFNγ (MCA1783PE, clone CC302, Bio-rad). Cell acquisition was performed on the BD LSRFortessa™ Cell Analyzer (BD Biosciences) and analyzed using FlowJo V10.6.2 (BD Biosciences).

### Statistical analysis

All data were analyzed with GraphPad Prism 9.1.0 software (GraphPad Software, Inc). Normality tests were employed to define the statistical test for the different data. Numerical values representing absolute or relative numbers are presented by means and standard deviation or by medians and interquartile range according to Gaussian distribution. Comparisons between two groups were performed using Mann-Whitney or Student’s t-test. Comparisons between multiple groups were performed using one-way ANOVA or the Kruskal-Wallis test. Multiple testing was corrected with Tukey’s multiple comparison test. Total area under the curve was calculated for body temperature measures across the groups. Fisher’s exact test was employed to analyze statistical significance for categorical data of clinical signs. The threshold for statistical significance was set at p<0.05.

## Results

### SARS-CoV-2 infection in ferrets

To evaluate the impact of a previous SARS-CoV-2 infection on a sequential influenza infection during the post-acute phase, we first studied viral replication and clinical signs in two groups of male ferrets that were infected with SARS-CoV-2 beta VOC during a 4 week period. In addition, we analyzed the humoral and cellular immune responses and sacrificed one of the two groups at day 28 to determine the pathological status of the respiratory tract at the day prior to the sequential influenza infection. The study was then continued with an influenza infection of naïve ferrets and the remaining group of ferrets that were infected with SARS-CoV-2 beta 4 weeks earlier (figure 1a). A control group consisted of mock-infected ferrets.

SARS-CoV-2 viral RNA was detected in nose and throat swabs of infected ferrets from 3 d.p.i. until at least 9 d.p.i., but not in mock-infected ferrets (figure 1b). SARS-CoV-2 virus copies were no longer detected at 29 d.p.i. The virus infected cells in both nose and throat, as evidenced by the detection of subgenomic viral RNA in swab samples (figure 1c). However, a productive infection was barely established, since low or sometimes absent viral titers were observed by TCID_50_ determination in nose swabs (peak at day 5, 1.6 ± 0.2 logTCID_50_/mL) and throat swabs (peak at day 3, 1.8 ± 0.2 logTCID_50_/mL) (figure 1d). Viral RNA was not detected in trachea or lung at day 28, but low copy numbers were found in the nasal turbinates of two out of six ferrets (figure 1e).

The clinical signs observed in SARS-CoV-2-infected ferrets were mild and of low frequency. Between 1 and 5 days after SARS-CoV-2 infection, increased breathing rhythm and/or decreased activity were observed in six out of twelve ferrets. Except for one ferret that showed mild decreased activity at 9 d.p.i. and had not shown any other clinical signs before, all clinical signs were absent at 6 d.p.i. (data not shown). SARS-CoV-2 infected ferrets did not present notable body weight loss in comparison to mock-infected ferrets (figure 2a), nor did they develop fever, as shown by normal daily variations in body temperature (figure 2b). Upon euthanasia, the ferrets did not show signs of COVID-19 in gross lung pathology and the relative lung weight, an indication of presence of edema secondary to inflammation in the lungs, was not increased (figure 2c). Rare histopathology findings in the lungs (figure 2d), together with moderate rhinitis in the nasal turbinates (figure 2e), reflect an absent to slight post-acute pathology of respiratory organs 28 days after SARS-CoV-2 infection.

**Figure 2.**
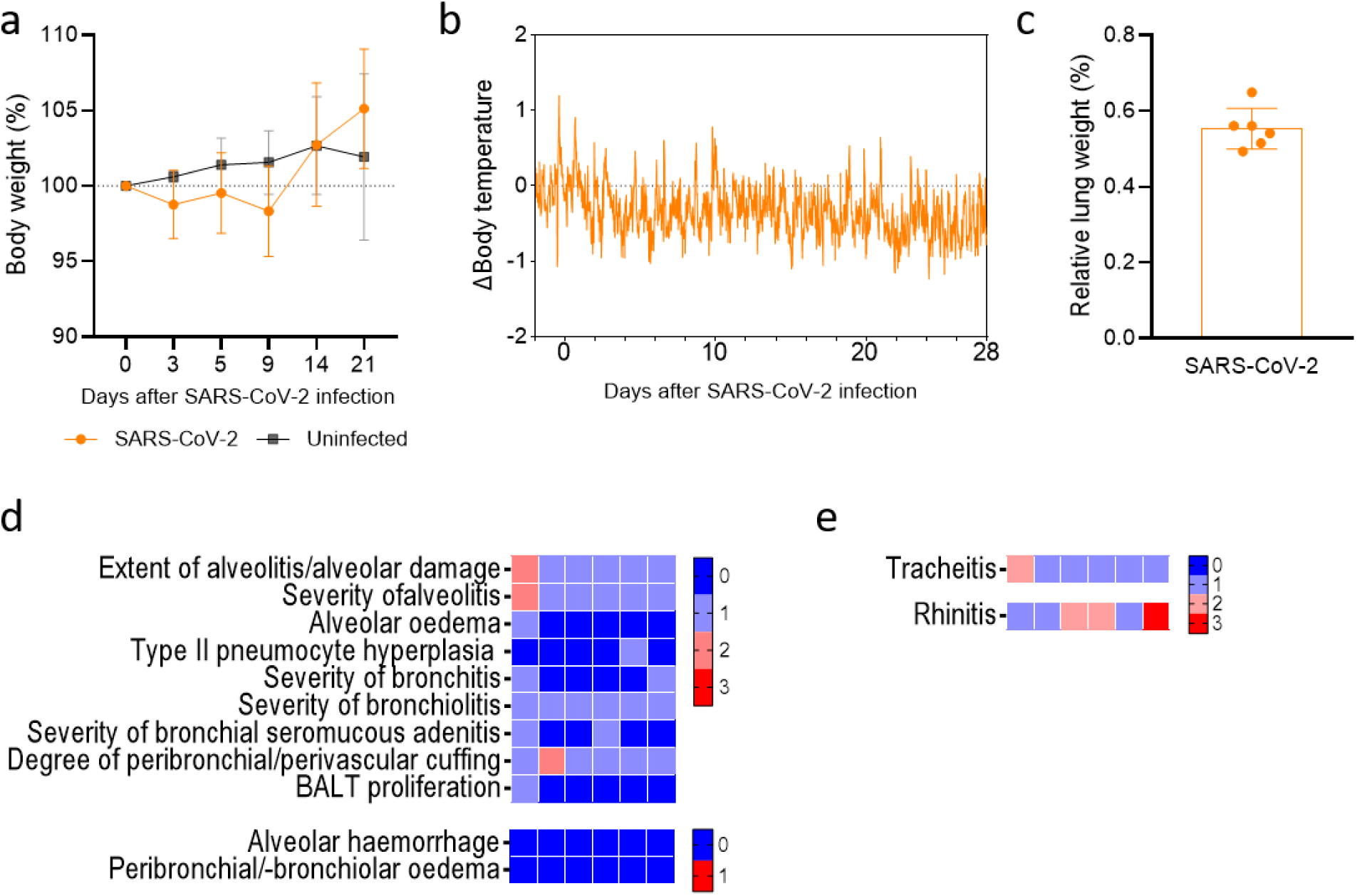
Clinical signs and histopathology in post-acute SARS-CoV-2 (B.1.351, beta VOC) experimental model. **(a)** Percentual body weight variation relative to baseline values (day 0) of beta VOC infected ferrets (n=12, orange dots) and mock-infected ferrets (n=3, gray squares) on 0, 3, 5, 9, 14 and 21 days post-infection (d.p.i.). **(b)** Differences (Δ) in body temperature up to 28 d.p.i. of beta VOC infected ferrets (n=5) relative to the baseline measurements recorded over a period of five days prior to infection. **(c)** Relative lung weight (RLW, %) of the body weight upon euthanasia on 28 d.p.i. of beta VOC-infected ferrets (n=6). The RLW is comparable to RLW of mock-infected ferrets (fig 4d). **(d-e)** Categorical heatmaps of histopathology represented in semi-quantitative intensity score, with color shading from blue (0) to red (3), or qualitative score of absence (0, blue) or presence (1, red), where indicated. **(d)** Lung histopathology and **(e)** histopathology scoring of inflammation of trachea (tracheitis) and nasal turbinates (rhinitis) of six SARS-CoV-2-infected ferrets. Data is visualized as mean with standard deviation and dots represent individual observations. For reference, data of body temperature and gross pathology and histopathology of mock-infected ferrets can be consulted in figures 4 and 5.

### Immune response in SARS-CoV-2 infection in ferrets

Despite the low level of replication of SARS-CoV-2 beta VOC (figure 1b-e), the ferrets did develop cellular immune responses as shown by restimulation of PBMCs with SARS-CoV-2 peptide pools and live virus in IFNγ-ELISpot (figure 3a). IFNγ production was observed in SARS-CoV-2 beta VOC-infected ferrets against both live alpha and beta VOCs, reflecting the similarity between these variants. This was sustained by the finding on IFNγ responses upon stimulation with peptide pools of spike protein (S) from the original SARS-CoV-2 strain. Additionally, we found an IFNγ response upon stimulation with nucleocapsid protein (N), but not with non-structural protein 12 (nsp12) which contains the RNA-dependent RNA polymerase (RdRp). No responses were detected in mock-infected ferrets, or in PBMCs stimulated with negative controls (mumps virus and HIV peptide pool). In addition to the ELISpot assay, we investigated the phenotype of the IFNγ-producing T cells by flow cytometry at 28 d.p.i. (figure 3b-c). PBMC stimulation with SARS-CoV-2 beta VOC live virus elicited clear IFNγ responses in both CD4 and CD8 T cells. No responses were detected after stimulation with peptide pools spanning for S protein (amino acid ranges 1-158 and 159-315) in CD8 T-cells, as the response was not distinguishable from the background (medium) and negative control (HIV peptide pool). For CD4 T cells, IFNγ response was distinctively found against the second portion (159-315) of the S protein. Antibody responses against S and RBD of SARS-CoV-2 beta VOC were found from 14 d.p.i. and reached a plateau on 21-28 d.p.i. (figure 3d).

**Figure 3.**
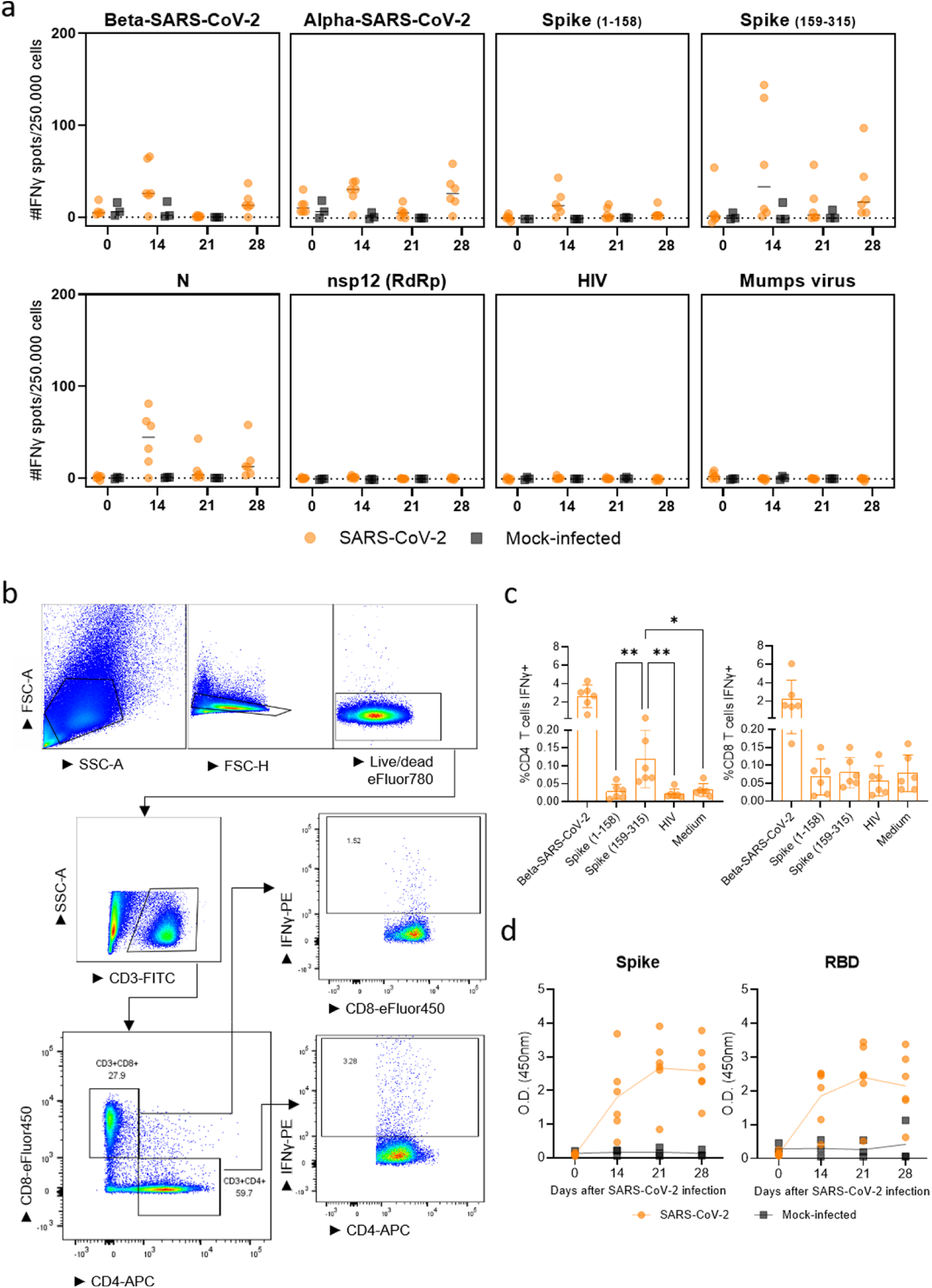
Cellular and humoral responses in SARS-CoV-2 (B.1.351, beta VOC) experimental model. **(a)** Cellular responses in PBMC on 0, 14, 21 and 28 days post-infection (d.p.i.) measured by IFNγ ELISpot against SARS-CoV-2 alpha VOC (with a culture-acquired mutation in the ORF1a region) and beta VOC, and SARS-CoV-2 peptide pools spanning spike protein (amino acid ranges 1-158 and 159-315), nucleocapsid protein (N) and RNA-dependent RNA polymerase (RdRp) – non-structural protein 12 (nsp12). Stimulations with mumps virus and HIV peptide pool were used as negative controls for virus and peptide pool stimulations, respectively. **(b)** Gating strategy of immunophenotyping of IFNγ-producing CD3+CD4+ and CD3+CD8+ T cells by flow cytometry and **(c)** relative (%) quantification of IFNγ-producing T cells on 28 d.p.i. The mock-infected group was not sampled due to its continuation in the study to serve as a control group for the subsequent H1N1 infection. **(d)** Humoral responses measured by ELISA assay. Optical density (O.D.) values evidence antibody responses against spike and receptor binding domain proteins of beta VOC on 0, 14, 21 and 28 d.p.i. Data is visualized as median (horizontal line, a) or mean (bar or connecting line, c and d) with standard deviation and dots represent individual observations. *p=0.01, **p<0.007, ANOVA and Tukey’s multiple comparison test.

### Influenza H1N1 infection and disease in ferrets with previous SARS-CoV-2 exposure

We next investigated if influenza virus-induced pathology and disease are enhanced during the post-acute phase of SARS-CoV-2 (SARS-CoV-2/H1N1 group) in comparison to H1N1 influenza virus infection alone (H1N1 group, figure 1a). H1N1 viral load and replication were assessed on days 0, 2, 4 and 5 d.p.i. with H1N1 influenza virus by TCID_50_ and RT-qPCR (figure 4a-b). Viral RNA and virus replication were observed as early as 2 d.p.i., in both nose and throat swabs. Upon euthanasia on 5 d.p.i., H1N1 RNA was found in lung, trachea and in nasal turbinates, while viable virus was only recovered from nasal turbinates by TCID_50_ assay (figure 4c). Overall, viral load and viral replication were similar between groups, irrespective of prior SARS-CoV-2 infection, and absent in mock-infected ferrets. Gross pathology showed signs of inflammation in the lungs of all animals infected with H1N1 and was absent in mock-infected ferrets (data not shown). Relative lung weight, representing edematous lung, was increased in all H1N1-infected ferrets compared to mock-infected ferrets, irrespective of previous SARS-CoV-2 infection (figure 4d). All H1N1-infected ferrets lost 6-8% of body weight from day 3 of H1N1 infection, which was not significantly different between groups (figure 4e). Increased body temperature in the first three days of H1N1 infection was observed in a similar curve between the groups (figure 4e). Altogether, viral burden and gross pathology did not indicate that influenza disease severity increased upon prior exposure to SARS-CoV-2. Nonetheless, clinical signs characteristic for influenza disease were more persistent in ferrets that received an H1N1 influenza virus infection consecutive to a SARS-CoV-2 infection (figure 4g). The relative frequency of decreased activity was similar in both groups of H1N1-infected ferrets on day 4. Labored breathing was observed in all H1N1-infected ferrets with previous SARS-CoV- 2 exposure (six out of six ferrets, 100%) in comparison to four out of five ferrets (80%) ferrets exposed to H1N1 only on day 4. On day 5, clinical signs were only observed in sequentially infected ferrets (SARS-CoV-2/H1N1), with more pronounced labored breathing (five out of six ferrets, 83%) and decreased behavioral activity (two out of six ferrets, 33%) while H1N1- infected ferrets without SARS-CoV-2 prior exposure were free of clinical signs.

**Figure 4.**
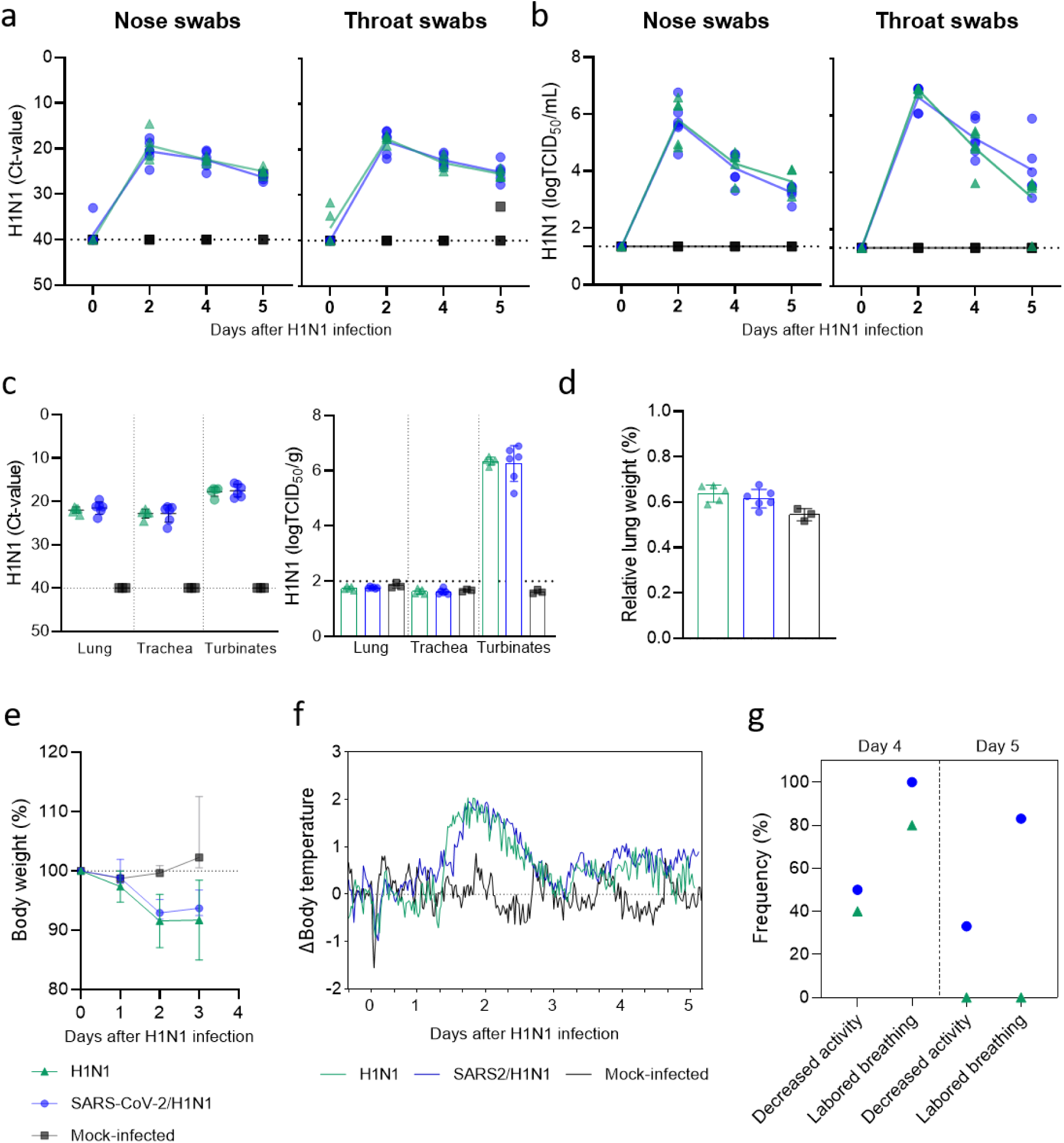
H1N1 influenza virus infection and disease in ferrets previously exposed to SARS-CoV-2 (B.1.351, beta VOC). H1N1 influenza viral load, virus replication and clinical disease in ferrets infected with H1N1 influenza virus only (H1N1, n=5, green triangles), in ferrets previously infected with SARS-CoV-2 beta VOC (SARS-CoV-2/H1N1, n=6, blue dots), and in mock-infected ferrets (n=3, black squares). **(a)** H1N1 viral RNA detected by RT-qPCR and **(b)** H1N1 virus replication detected by TCID_50_ assay in nose and throat swabs on 0, 2, 4 and 5 days post infection (d.p.i.). **(c)** H1N1 viral RNA detected by RT-PCR and viral replication assessed by TCID_50_ in lung, trachea and nasal turbinates on 5 d.p.i. **(a-c)** Horizontal dotted lines depict the limit of detection by RT-PCR and TCID_50_ assay. **(d)** Relative lung weight (%) of the body weight upon euthanasia on 5 d.p.i.. **(e)** Percentual body weight variation relative to weight on the day of infection (day 0) of H1N1- or mock-infected ferrets on 0, 2, 4 and 5 d.p.i. **(f)** Differences (Δ) in body temperature up to 5 d.p.i. relative to the baseline measurements recorded five days prior to infection. **(g)** Recorded clinical signs of influenza disease of decreased activity and labored breathing. Percentage of ferrets that developed the cited clinical signs in the different groups on 4 and 5 d.p.i. Data is visualized as mean and datapoints represent individual observations (a-d). Bars are set at the mean and standard deviation is shown (d and e).

### Pathology of H1N1 infection and SARS-CoV-2 in respiratory organs of ferrets

Because prior exposure to SARS-CoV-2 appeared to marginally worsen influenza disease symptoms, we wondered if differential pathology could be observed in the respiratory tract. In general, moderate to severe broncho-interstitial pneumonia, bronchoadenitis and thickening of the alveolar septa and large amounts of oedema in the alveolar lumina characteristic for influenza disease were observed (figure 5a). Severe rhinitis was observed in all H1N1-infected ferrets (figure 5b), which was also present in moderate intensity in SARS-CoV-2-infected ferrets, long after the SARS-CoV-2 infection was cleared (figure 2e). Mild to moderate tracheitis was observed (figure 5b). Regardless of prior SARS-CoV-2 exposure, the histopathology findings were homogenous between the groups, except for a trend of increased type II pneumocyte hyperplasia especially in the cranial lobe of the lungs and moderate bronchitis found more prominent in the caudal lobe in the SARS-CoV-2 group sequentially infected with H1N1 (figures 5a and 5c). Altogether, these results suggest that a previous mild SARS-CoV-2 beta VOC infection does not notably enhance the respiratory tract pathology induced by a sequential infection with influenza.

**Figure 5.**
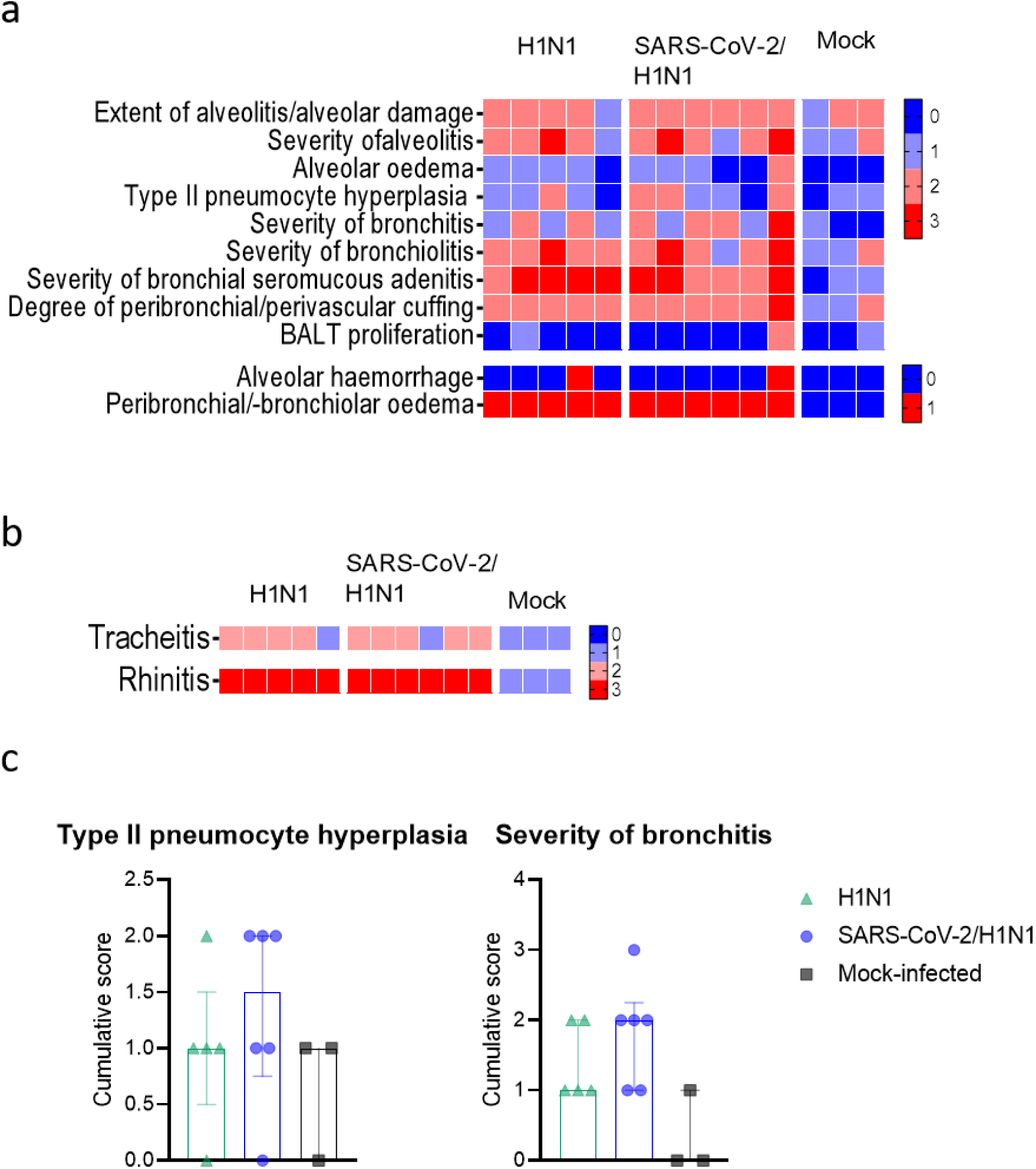
Histopathology of H1N1 infection in ferrets previously exposed to SARS-CoV- 2 (B.1.351, beta VOC). Categorical heatmaps of histopathology represented in semi- quantitative intensity score, with color shading from blue (0) to red (3), or qualitative score of absence (0, blue) or presence (1, red), where indicated. **(a)** Lung histopathology and **(b)** Histopathology scoring of inflammation of trachea (tracheitis) and nasal turbinates (rhinitis). **(c)** Type II pneumocyte hyperplasia and severity of bronchitis in ferrets infected with H1N1 influenza virus only (H1N1, n=5, green triangles), or in ferrets previously infected with SARS-CoV-2 (SARS-CoV-2/H1N1, n=6, blue dots), and mock-infected ferrets (n=3, black squares). Data presented as median and interquartile range.

## Discussion

In this study, we investigated if a previous exposure to SARS-CoV-2 enhances influenza disease during the post-acute phase of SARS-CoV-2 infection. Following a resolved SARS-CoV-2 infection, ferrets showed a tendency of increased influenza clinical symptoms by a sequential H1N1 infection, compared to H1N1 infection without prior exposure to SARS-CoV- 2 and only minor effects were found in histopathology. Resolved histopathology of the respiratory organs 4 weeks after infection with the beta variant of SARS-CoV-2, following a mild viral replication, may have only marginally affected a sequential H1N1 infection. It remains to be determined if a robust SARS-CoV-2 infection would have a greater impact on the sequential H1N1 infection, as we found that a mild infection already has a slight negative impact on the influenza clinical disease.

Ferrets are a suitable model of COVID-19 for features such as viral replication kinetics, virus dissemination and transmission compared to other small animal models [28, 29]. Clinical presentation is absent or mild, and histopathology findings usually show variable grades of inflammatory infiltration in the upper and/or lower respiratory tract, which is not homogeneous among reports [22, 30, 31]. However, a large share of these studies relate to the original strains of SARS-CoV-2, and little is yet known about the susceptibility of the ferret to VOCs. While it has been shown that the SARS-CoV-2 alpha VOC successfully replicates in ferrets [32, 33], the absence of productive infection with beta VOC in ferrets was recently reported [33]. To our knowledge, only one investigation reported successful replication with the beta-SARS-CoV-2 variant in female ferrets, with a 100-times lower dose than that we used in this study [34]. Using a high infectious dose (10_7_ TCID_50_/mL) in male ferrets, we found that the beta VOC replicated only to low levels, despite a successful infection as evidenced by the detection of subgenomic viral RNA. In another study, beta VOC was outcompeted by co-infection with alpha VOC in ferrets and thus proves less fit in ferrets [32]. The emergence of the beta VOC is believed to have resulted from immune escape in a partly immunized population, whereas in a naive population it proved less fit than the alpha variant [32, 35], which may reflect the low replication found in naïve ferrets in this study.

Regardless of the low level of productive infection of beta VOC, cellular and humoral responses were induced. IFNγ production was found in PBMCs upon stimulation with other SARS-CoV-2 strains, indicating that the mutations that appeared in the VOCs did not affect the cellular response. In fact, using peptide stimulations based on the original strain, N induced IFNγ production, but a more prominent response was found against the S protein. In line with our results, immune responses against S and N proteins are described for both ferrets and humans [23, 36, 37], while nsp12 (RdRp) was shown to induce a low response [38]. SARS-CoV-2-specific cellular and humoral responses were associated with protective immunity in human COVID-19 [39], which may have also contributed with the controlled SARS-CoV-2 infection in ferrets.

Prolonged clinical symptoms of COVID-19 are considered a syndrome after long resolved SARS-CoV-2 infection in humans known as post-acute COVID-19 or long-COVID [8, 9]. The knowledge of the lung pathology in post-acute COVID-19 cases is scarce, as most information on human lung pathology is known from acute fatal cases [40]. In a previous study using SARS-CoV-2 with the D614G mutation, we observed inflammatory infiltration in the upper respiratory tract and hyperplasia of bronchus-associated lymphoid tissue in the lungs of infected male ferrets on 21 d.p.i. [23]. The late histopathology findings may reflect, in part, the extended signs and symptoms observed in human long-COVID. Studies performing intranasal infection with the original strain of SARS-CoV-2 showed pathology alterations in the respiratory tract on 21 d.p.i., after the virus was cleared, suggesting an extended effect of the infection [23, 29]. Also at 21 d.p.i., female ferrets were shown to develop mild (peri)bronchiole and interalveolar septal infiltration [29]. In the current study, we looked at these effects 28 d.p.i., as we expected that the histopathology found in other studies at 21 d.p.i. would likely not yet be resolved. Yet, only a moderate inflammation in the nasal turbinates was observed, and no histopathology alterations were found in the lower respiratory tract. We assume, however, that the absence of pathology in the present study is due to the low level of productive infection when using the beta VOC, since effective SARS-CoV-2 viral replication frequently resulted in pathology manifestation in ferrets in other studies [19, 22, 23, 29, 30, 41].

It has been shown that co-infection between influenza and SARS-CoV-2 results in more severe disease in animal models through enhanced lung damage [16, 17, 19, 42]. This simultaneous infection was also reported in human patients [10, 11, 43]. In our study, we looked at the effects of a sequential influenza infection during the post-acute phase of SARS-CoV-2 (28 days), as an influenza infection is less likely to happen simultaneously or shortly after a SARS-CoV-2 infection, as compared to an infection during the time span in which humans suffer from long-COVID. In the current study, we established a reproductive H1N1 influenza virus infection, with clinical signs, viral replication kinetics and histopathology characteristic for an infection with this subtype [44]. The intensity and number of histopathological observations in the respiratory tract, however, was not clearly different between ferrets that were previously infected with SARS-CoV-2 and ferrets that were not. Nevertheless, a trend for a more severe bronchitis and type II pneumocyte hyperplasia score was observed in ferrets that were sequentially infected with SARS-CoV-2 and then H1N1 influenza virus, although the difference was not statistically significant. This observation could explain the tendency of more persistent alterations in clinically observed respiratory function in the sequentially infected ferrets. Both SARS-CoV-2 and influenza viruses infect type II pneumocytes [45, 46], which could account for the hyperplasia in response to the infection in both occasions, as well as an indication that acute lung injury caused by influenza could be worsened by a previous SARS-CoV-2 infection.

In conclusion, our data suggest that there is likely a SARS-CoV-2 variant-dependent effect on the severity of sequential influenza infection. Because a slight enhancement of influenza disease was observed even upon a mild preceding SARS-CoV-2 infection, further studies are necessary to confirm the impact of more virulent SARS-CoV-2 VOCs and the development to long-COVID. Currently, patients suffering from long-COVID are not specifically considered as a risk group for influenza disease. In light of the results found herein, despite being subtle, it may be advisable to preemptively include this group in the influenza vaccination campaign.

## Acknowledgements

We would like to thank Helena Pinheiro Guimarães for her help during the animal experiments, to Maarten Emmelot and Gabriel Goderski for providing the mumps and SARS-CoV-2 viruses and to Dr. Elena Pinelli Ortiz and Dr. Willem Luytjes for their critical review of the manuscript.

## Supplementary material

**Table 1.** Baseline serology of the ferrets included in this study.

| Group | CHV | FCOV | CCV | Aleutian | Reference |
| --- | --- | --- | --- | --- | --- |
| **SARS-CoV-2** | 150 | 150 | 450 | <100 | ≤100 |
|  | 100 | 150 | 800 | <100 | ≤100 |
|  | 100 | 200 | >800 | <100 | ≤100 |
|  | 150 | 250 | >800 | <100 | ≤100 |
|  | 100 | 200 | >800 | <100 | ≤100 |
|  | 100 | ≤100 | 225 | <100 | ≤100 |
| **H1N1** | <100 | <100 | 225 | <100 | ≤100 |
|  | 300 | 150 | 500 | 100 | ≤100 |
|  | 300 | 150 | 300 | 200 | ≤100 |
|  | 150 | 100 | 300 | ≤100 | ≤100 |
| **SARS-CoV-2/H1N1** | 100 | ≤100 | 100 | 100 | ≤100 |
|  | 100 | 150 | 325 | <100 | ≤100 |
|  | 150 | 150 | 325 | <100 | ≤100 |
|  | 150 | 200 | 175 | <100 | ≤100 |
|  | 250 | 400 | 500 | 200 | ≤100 |
|  | 100 | <100 | 200 | <100 | ≤100 |
|  | 150 | <100 | 750 | 100 | ≤100 |
|  | ≤100 | 100 | 150 | <100 | ≤100 |
|  | 100 | 200 | 150 | <100 | ≤100 |
|  | 150 | 100 | 350 | 100 | ≤100 |
|  | 150 | <100 | 650 | <100 | ≤100 |
| **Mock-infected** | 100 | <100 | 100 | <100 | ≤100 |
|  | 150 | 100 | 150 | <100 | ≤100 |
|  | 200 | 150 | 650 | 100 | ≤100 |

Note: Data is expressed in optical density values obtained by functional ELISA assay from ferret sera for ferret coronavirus (FCOV), enteric coronavirus (CCV), Aleutian disease and canine herpes virus (CHV).

